# Azithromycin Plus Zinc Sulfate Rapidly and Synergistically Suppresses IκBα-Mediated In Vitro Human Airway Cell ACE2 Expression for SARS-CoV-2 Entry

**DOI:** 10.1101/2021.01.19.427206

**Authors:** Chia-Wei Chang, Ming-Cheng Lee, Bor-Ru Lin, Yen-Pei Lu, Yih-Jen Hsu, Chun-Yu Chuang, Tsung-Tao Huang, Yin-Kai Chen

## Abstract

Large-scale efforts have been persistently undertaken for medical prophylaxis and treatment of COVID-19 disasters worldwide. A variety of novel viral spike protein-targeted vaccine preparations have recently been clinically distributed based on accelerated approval. We revisited the early but inconclusive clinical interest in the combination of azithromycin and zinc sulfate repurposing with safety advantages. In vitro proof of concept was provided for rapid and synergistic suppression of ACE2 expression following treatments in human airway cells, Calu-3 and H322M. The two representative ACE2-expressing human airway cells indicate the upper and lower respiratory tracts. Prophylactic and early therapeutic roles of azithromycin combined with zinc are proposed for virus cellular entry prevention potential bridging to effective antibody production.

## Introduction

Cell surface angiotensin-converting enzymes 2 (ACE2) of the respiratory tract is a well-established critical entry of SARS-CoV-2 into infected cells [1–3]. ACE2 mRNA expression was shown to be reduced from tracheobronchial to bronchioloalveolar regions [4]. ACE2 expression was upregulated following viral infection [5, 6], interferon exposure [5, 6] and smoking [5, 6]. It has been postulated that hyperactivation of the transcription factor NF-κB following ACE2-mediated viral entry, most likely in nonimmune cells, including lung epithelial cells, resulted in cytokine release syndrome [7]. NF-κB is a major transcription factor that regulates the genes responsible for innate and adaptive immune responses. Human type II pneumocytes (AT2) are one of the primary targets for SARS-CoV-2 infection [6]. It was shown in an induced pluripotent stem cell-derived AT2 model that NF-κB signaling was rapidly and persistently upregulated upon SARS-CoV-2 infection [8]. Azithromycin (AZT), a second-generation macrolide with broad spectrum antibacterial activity, has drawn early clinical attention in hot drug repurposing for the recurrence of COVID-19 patients. [9–12] In addition to its antimicrobial activity resulting from bacterial protein synthesis inhibition [13], AZT protects against viral entry into A549 lung cancer cell lines [14] and viral infections of airway epithelial cells through reduced viral replication and increased interferon responses [15–17]. AZT was demonstrated in vivo to suppress NF-κB activation and concomitant pulmonary inflammation [18]. TNF-α-induced NF-κB DNA binding activity, IκBα degradation and IL-6/IL-8 release in tracheal cells of human origin were shown to be dose-dependently inhibited in vitro following AZT [19].

Zinc is a trace element supplement with clinical benefits in respiratory tract infections [20]. Zinc deficiency prevalence of 26 % was ever reported in a case-control study of adults aged 50 years or older visiting an Ohio outpatient clinic between 2014 and 2017 [21]. Patients with COVID-19 had significantly lower serum zinc levels than normal controls [22]. More complications are developed in COVID-19 patients with zinc deficiency [22]. The prophylactic and therapeutic roles of zinc supplementation are currently under investigation. [23, 24] Prevention of viral entry has been postulated to be a potential mechanism for zinc antiviral actions [20]. Increased NF-κB DNA binding activity and IκBα mRNA expression were reported in lungs from a septic mouse model with zinc deficiency [25].

Short-course AZT has long been clinically used for atypical pneumonia cases with excellent safety and tolerability. Zinc supplementation is a relatively popular companion with vitamin complexes. Here, we showed markedly synergistic ACE2 suppression in two distinct ACE2-expressing human airway cells by combined AZT and zinc treatments. Accordingly, we provided laboratory support to repurpose AZT plus zinc for the early clinical intervention of COVID-19.

## Results and Discussion

Because airway is the primary and lethal target involved in COVID-19, Calu-3, H322M, H522, H460, H1299, and A549 human airway cells were screened for endogenous ACE2 expression. Calu-3, widely used for COVID-19 studies, and H322M were selected for subsequent exploration because of their endogenous ACE2 expression (Fig. 1A). The Calu-3 cells generated from human proximal bronchial adenocarcinoma [26] are characterized by differentiated, functional human airway epithelial cells [27]. These cells were also proposed to be a suitable model for the human nasal mucosa [28]. Bronchoalveolar lavage analyses from COVID-19 patients disclosed aberrant macrophage and T cell responses [29] as well as bronchoalveolar immune hyperactivation [30]. Our drug repurposing for the critical bronchoalveolar involvement of COVID-19 was investigated with H322M, a bronchioalveolar cell line, in addition to Calu-3 for proximal airway involvement.

**Figure 1.**
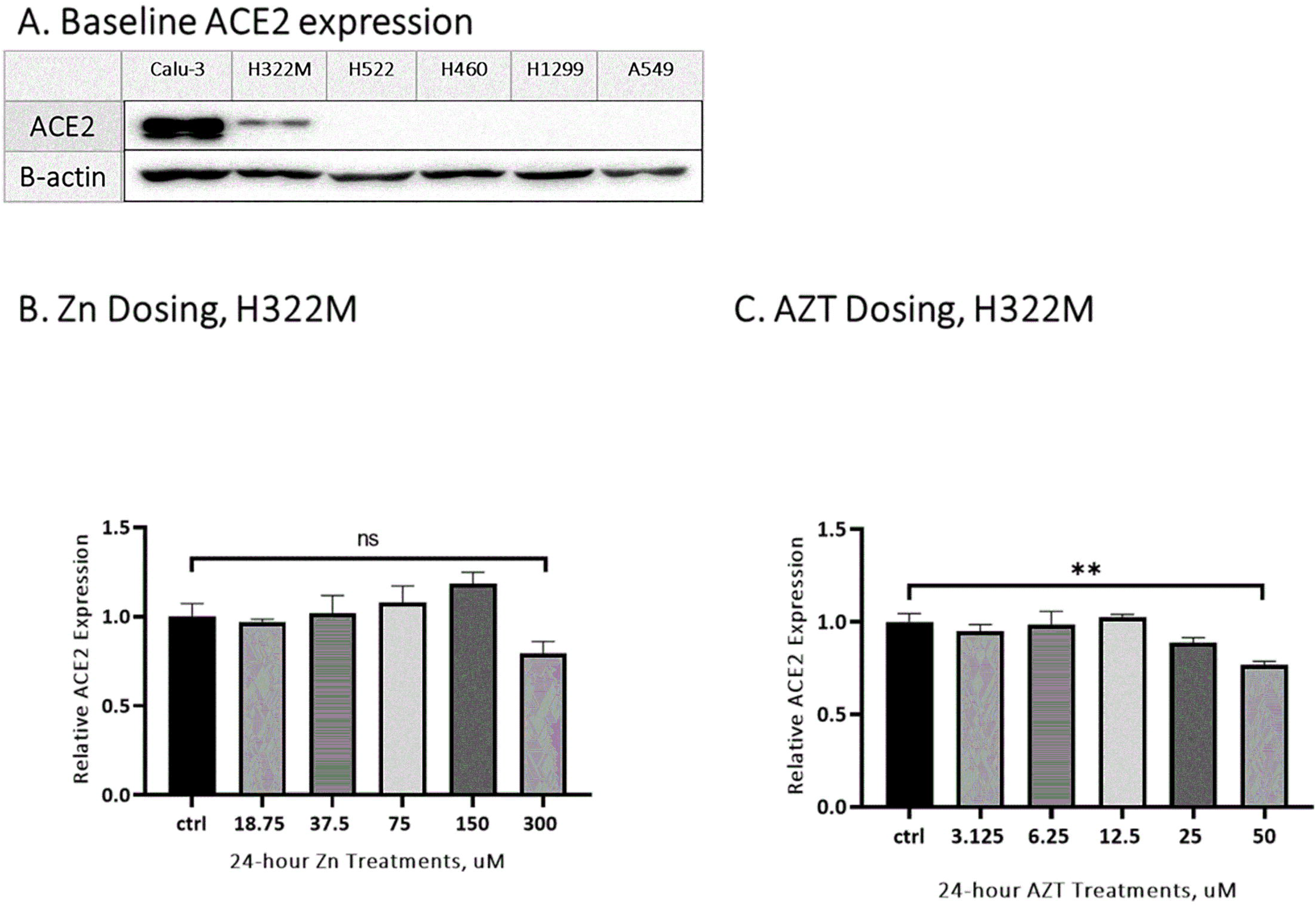
Azithromycin (AZT) but not ZnSO4 (Zn) treatment alone decreased endogenous ACE2 expression in the human lower airway H322M cells. (A) Endogenous ACE2 expression was screened by Western blot using beta-actin as a loading control. ACE2-expressing Calu-3 and H322M cell lines were selected for subsequent exploration. (B) H322M cells were treated with serial concentrations of Zn (0, 18.75, 37.5, 75, 150 and 300 μM) for 24 hours, and total RNA was collected for *ACE2* quantitation by real-time qRT-PCR assays. The data were normalized to *GAPDH* expression and presented as the mean ± SEM (n = 3). *ACE2* expression was potentially decreased under 24-hour treatment with 300 μM Zn. This concentration was determined for combination treatment with AZT. (C) H322M cells were treated with a concentration series of AZT (0, 3.125, 3.25, 12.5, 25 and 50 μM) for 24 hours, and total RNA was collected for *ACE2* quantitation by real-time qRT-PCR assays. The data were normalized to *GAPDH* expression and presented as the mean ± SEM (n = 3). (**p = 0.001 to 0.01). *ACE2* expression was significantly decreased under 24-hour treatment with 50 μM AZT. This concentration was adopted for combination treatment with Zn.

Symptoms, including fever, dyspnea and hypoxia, were rapidly improved following high-dose zinc rescue with different preparations in a consecutive COVID-19 case series. [31] A randomized controlled trial for high-dose intravenous zinc was initiated as adjunctive therapy in SARS-CoV-2-positive critically ill patients [32]. H322M was therefore treated using 24-hour serial doses of 18.75, 37.5, 75, 150 and 300 μM ZnSO4 (Zn) (Fig. 1B). Zinc at 300 μM was determined for the subsequent drug repurposing combination study because ACE2 expression was potentially decreased following 24-hour treatment (Fig. 1B). 50 μM AZT induced significant in vitro anti-rhinoviral activities in normal primary bronchial epithelial cells [15] and those cells from children with cystic fibrosis [16]. Lower in vitro anti-rhinoviral levels of AZT were also reported for bronchial epithelial cells from patients with chronic obstructive lung disease [17]. Accordingly, H322M was treated with 24-hour serial doses of 50, 25, 12.5, 6.25 and 3.125 μM (Fig. 1C). An AZT concentration of 50 μM was chosen for the Zn combination because ACE2 expression was significantly decreased in H322M following 24-hour treatment (Fig. 1C).

This study’s most impressive finding was the rapidly suppressed endogenous ACE2 expression of H322M under 24-hour treatment of 300 μM Zn combined with 50 and 25 μM AZT (Figs. 2A and 3A). Compared to 50 and 25 μM AZT treatments alone, 300 μM Zn showed a significant synergistic suppressive effect on ACE2 expression in H322M (Figs. 2A and 3A). ACE2 expression in H322M was further decreased following 48-hour treatment with Zn and AZT combinations, especially 50 μM AZT (Fig. 2B). ACE2 mRNA expression of Calu-3 was significantly reduced following 24-hour treatment of 300 μM Zn alone and in combination with 50 and 25 μM AZT (Fig. 3C). The markedly suppressive effect on ACE2 protein expression in Calu-3 was found 24 hours later following the suppression of ACE2 mRNA expression. Similar to H322M, 48-hour combination treatment with 300 μM and 50 μM AZT showed the most suppressive effect on ACE2 expression in Calu-3 (Fig. 2D).

**Figure 2.**
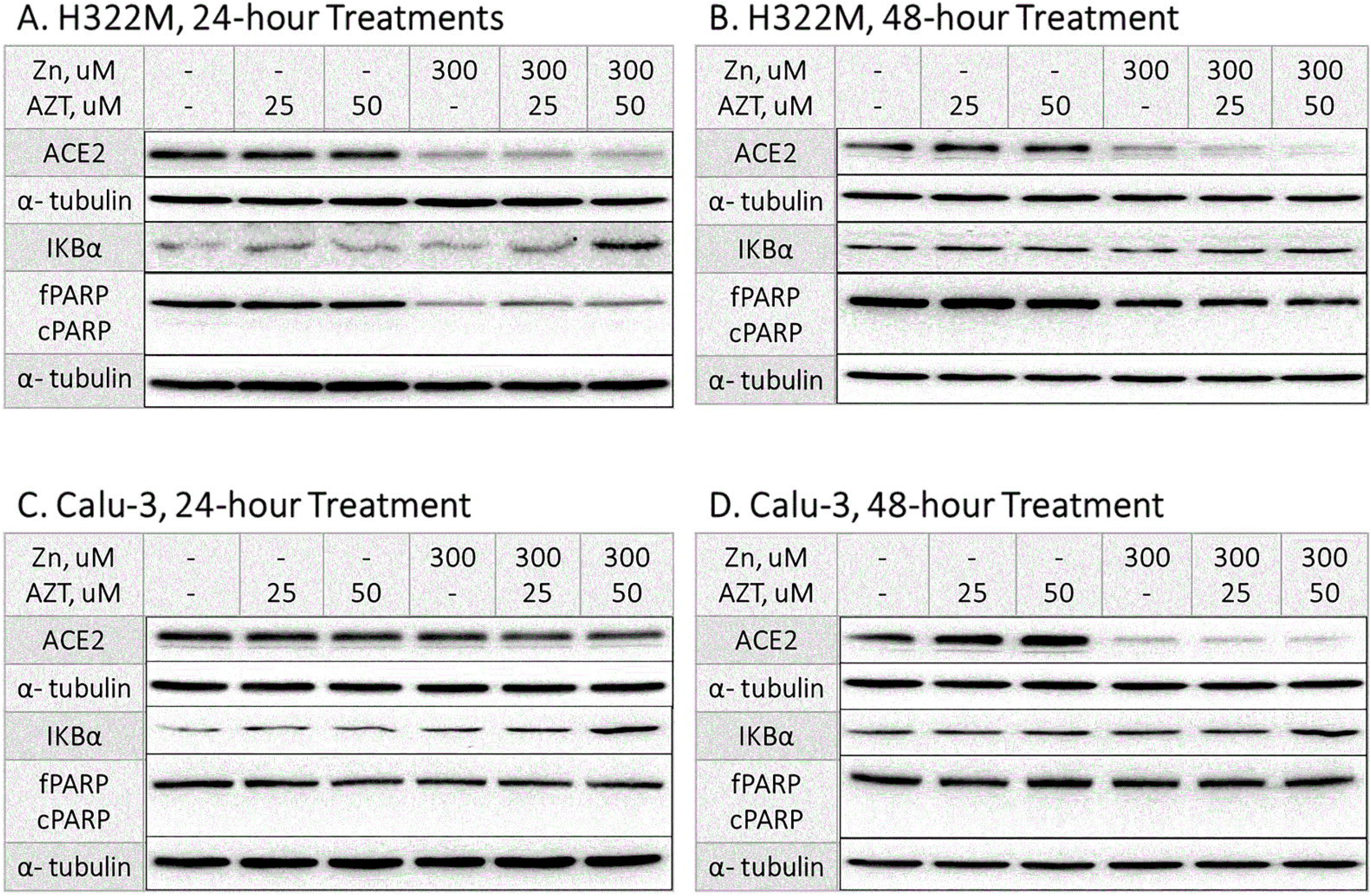
Endogenous ACE2 expression in H322M and Calu-3 cells was markedly suppressed by Zn and in combination with AZT in a time- and dose-dependent manner. (A and B) H322M cells were treated with 25, 50 μM AZT and 300 μM Zn in combination for 24 (A) and 48 (B) hours. Treated cells were lysed for Western blot analysis of ACE2 and PARP with α-actin as a loading control. ACE2 expression was suppressed without detectable cleaved PARP by Zn alone and synergistically with AZT in a time- and dose-dependent manner. IKB-α expression was obviously increased following 24 hours of treatment with 300 μM Zn combined with 50 μM AZT. (C and D) Calu-3 cells were treated with 25, 50 μM AZT and 300 μM Zn in combination for 24 (C) and 48 (D) hours. Compared to H322M treated in the same ways, ACE2 expression was suppressed without detectable cleaved PARP by Zn alone and synergistically with AZT in a more time-dependent manner. Similarly, IKB-α expression was obviously increased following 24 hours of treatment with 300 μM Zn combined with 50 μM AZT.

**Figure 3.**
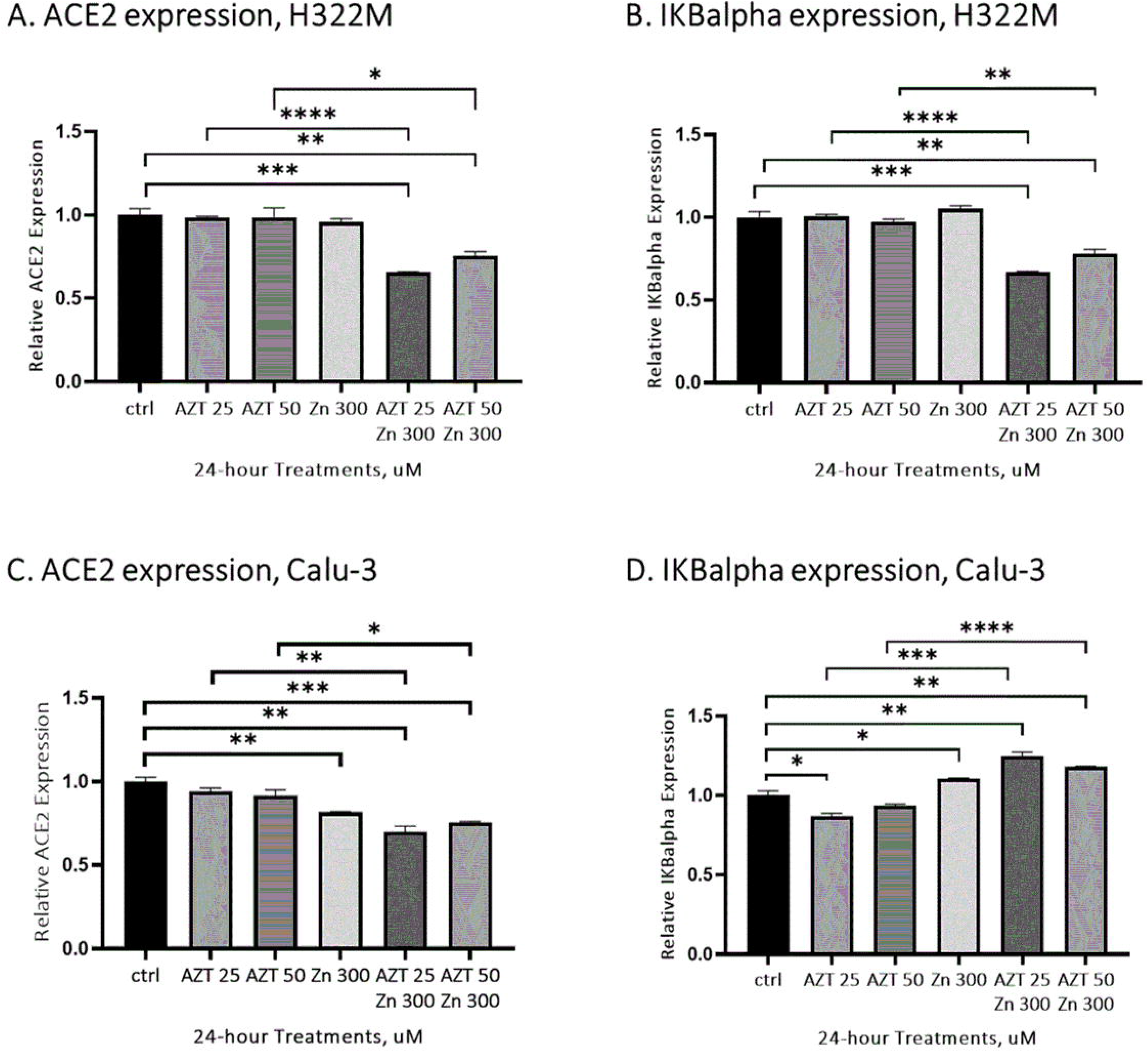
Endogenous ACE2 and IKBα expression was synergistically regulated by combined treatments of Zn and AZT. (A and B) H322M cells were treated with 25, 50 μM AZT and 300 μM Zn in combination for 24 hours. Total RNA was collected for *ACE2* (A) and *IKB*α (B) quantitation by real-time qRT-PCR assays. The data were normalized to GAPDH expression and presented as the mean ± SEM (n = 3). (*p = 0.01 to 0.05, ** p = 0.001 to 0.01, ***p = 0.0001 to 0.001 and ****p = < 0.0001). Compared to the control and AZT alone, 300 μM Zn synergistically decreased *ACE2* and *IKB*α expression with 25 and 50 μM AZT treatments, respectively. (C and D) Calu-3 cells were treated with 25, 50 μM AZT and 300 μM Zn in combination for 24 hours. Total RNA was collected for *ACE2* (A) and *IKB*α (B) quantitation by real-time qRT-PCR assays. The data were normalized to GAPDH/RPLP0 expression for H322M/Calu-3 cells respectively and presented as the mean ± SEM (n = 3). (*p = 0.01 to 0.05, ** p = 0.001 to 0.01, ***p = 0.0001 to 0.001 and ****p = < 0.0001). Zn showed a similar synergistic suppressive effect on *ACE2* expression in AZT treated Calu-3 cells. Conversely, *IKB*α expression was synergistically upregulated following 25 and 50 μM AZT treatments combined with 300 μM Zn compared to AZT alone.

Certain NF-κB dimeric transcription factors and their activities are tightly repressed by three inhibitors, IκBα, IκBβ and IκB◻, through the formation of stable IκB–NF-κB complexes. It was demonstrated that rapid degradation of free IκBα is critical for NF-κB activation [33]. Our H322M-associated results indicated that Zn and AZT combination treatments altered IκBα degradation and contributed to rapid repression of endogenous ACE2 (Figs. 2A, 2B, 3A and 3B). Such synergistic effects were most prominent following 24-hour treatment with 300 μM Zn and 50 μM AZT (Figs. 2A and 2B). In contrast to the synergistic effect on IκBα degradation in H322M, IκBα expression in Calu-3 was rapidly upregulated following 24-hour treatment of 300 μM Zn combined with 50 and 25 μM AZT (Figs. 2C and 3D). A similar synergistic ACE2 suppressive effect was most prominent following 48-hour 300 μM Zn treatment combined with 50 μM AZT (Figs. 2C and 2D). Accordingly, the underlying mechanisms of IκBα involved in ACE2 repression might be cell type specific and worth further investigation.

Membrane-tethered MUC1 belongs to one of the major components of mucus [34]. In vitro MUC1 overexpression was demonstrated to limit influenza A virus (IAV) infection [35]. In vivo MUC1 expression impeded IAV and reduced IAV disease severity [35]. NF-κB activation in vitro via toll-like receptor (TLR) pathways, including TLR 2, 3, 4, 5, 7 and 9, was counteracted by the MUC1 overexpression [36]. There is increasing evidence and a therapeutic interest in TLR pathway modulations for COVID-19 [37–39]. Elevated MUC1 mucin protein levels were found in the airway mucus of critically ill COVID-19 patients [40]. Another interesting finding of this study was that MUC1 expression in H322M and Calu-3 cells was most significantly increased following 24 hour treatment with 300 μM Zn alone and a lesser degree after combined treatment with 50 and 25 μM AZT (Figs. 4A and 4B).

**Figure 4.**
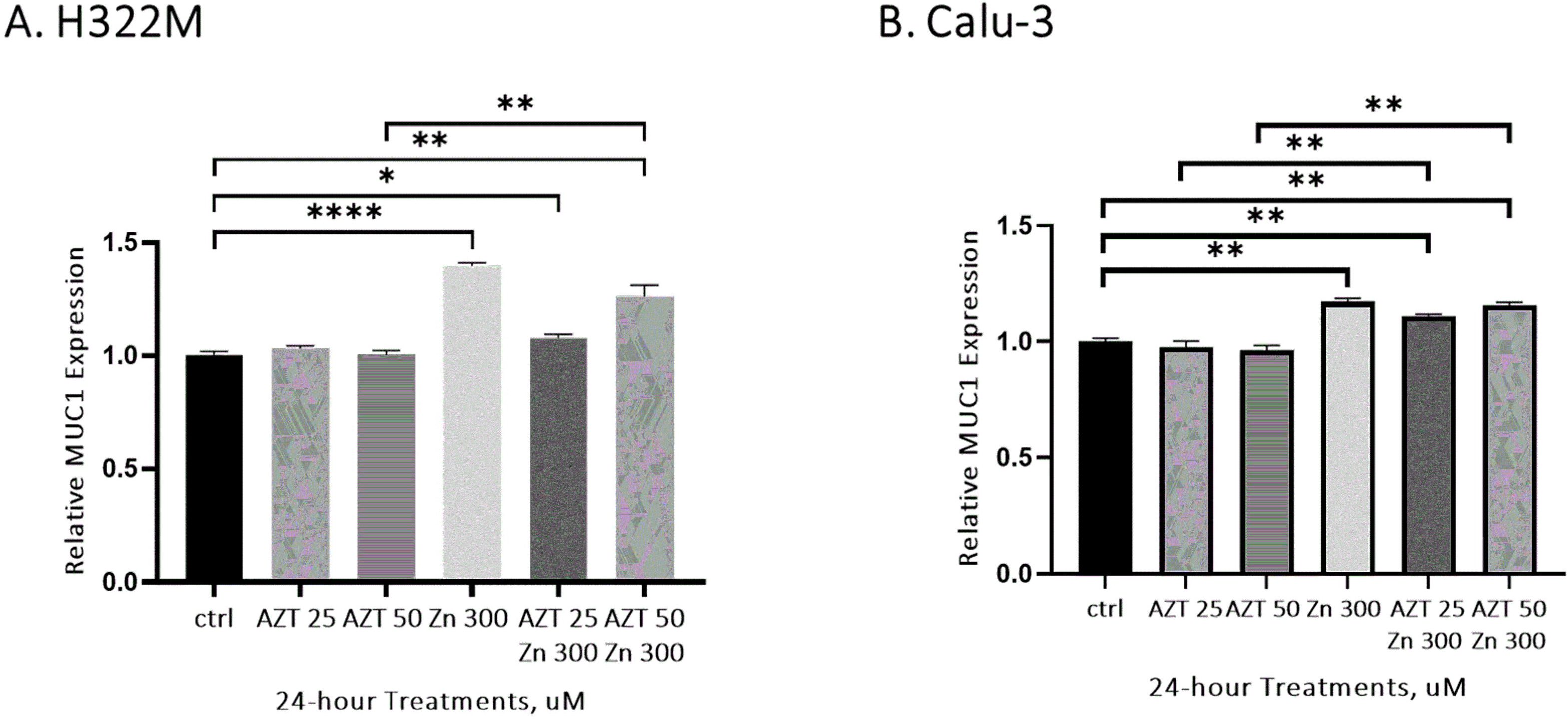
Endogenous MUC1 expression was significantly increased by Zn treatment alone and to a lesser degree in combination with AZT. H322M (A) and Calu-3 (B) cells were treated with 25, 50 μM AZT and 300 μM Zn in combination for 24 hours. Total RNA was collected for *MUC1* quantitation by real-time qRT-PCR assays. The data were normalized to GAPDH/RPLP0 expression for H322M/Calu-3 cells respectively and are presented as the mean ± SEM (n = 3). (*p = 0.01 to 0.05, ** p = 0.001 to 0.01, ***p = 0.0001 to 0.001 and ****p = < 0.0001). Compared to the control, *MUC1* expression was significantly increased following 300 μM Zn treatment alone and to a lesser degree combined with AZT in H322M and Calu-3 cells. In H322M, *MUC1* expression was synergistically increased following 50 μM AZT treatments combined with 300 μM Zn compared to AZT alone. In Calu-3 cells, *MUC1* expression was synergistically increased following 25 and 50 μM AZT treatments combined with 300 μM Zn compared to AZT alone.

The largest published retrospective case series reported that early outpatient treatment with zinc plus low-dose hydroxychloroquine and azithromycin significantly reduced the hospitalization rate [41]. Compared with the hydroxychloroquine-based standard of care, the addition of AZT did not show improved clinical outcomes in an open-label multicenter randomized clinical trial performed among patients with severe respiratory COVID-19 in Brazil [42]. Alternatively, large-scale randomized trials did not reveal hydroxychloroquine’s clinical benefits of in patients with COVID-19 [43, 44]. Accordingly, the feasible clinical timing and combination choice with AZT remain unelucidated.

A computed model of the AZT-Zn^++^ complex demonstrated its potential against the replication and assembly of SARS-CoV-2 particles. We are the first to present in vitro evidence that Zn combined with AZT rapidly and significantly suppresses endogenous ACE2 expression and increases MUC1 expression in Calu-3 and H322M cells, representative of the human upper and lower respiratory tracts, respectively. Both ACE2 suppression and increased MUC1 expression were consistently demonstrated in the two human airway cells, indicating the prophylactic and early therapeutic potential of AZT and Zn repurposing combination for COVID-19. With threats of mutations located on the viral spike protein [44], we demonstrated a potentially rapid bridging way before effective antibody production after vaccination by suppressing key cellular virus entry. The effective high-dose combination of Zn and AZT showed here could be translated into a loading dose modality in preclinical studies and clinical trials.

## Materials and Methods

### Cell lines and drug treatments

<u>The</u> H322M, H522, H460 and A549 human airway cells were courtesy of SLY (National Taiwan University), and the H1299 lung cancer cell line was courtesy of CCH (National Taiwan University Hospital). Calu-3 human upper airway cells were purchased from the American Type Culture Collection (ATCC) (Manassas, VA, USA). The Calu-3 cells were cultured with MEM (Thermo Fisher Scientific, Waltham, MA, USA) containing 20% (v/v) fetal bovine serum (Thermo Fisher Scientific), 100 mM sodium pyruvate, nonessential amino acids and penicillin-streptomycin in a humidified 5% CO_2_ atmosphere. The H322M cells were cultured with RPMI medium (Thermo Fisher Scientific) containing 10% (v/v) fetal bovine serum (Thermo Fisher Scientific) and penicillin-streptomycin at 37◻°C in a humidified 5% CO_2_ atmosphere.

After seeding at a density of 1×10^6^ cells per 6-well plate, Calu-3 and H322M cells were incubated for 20 h at 37 °C in 5% CO_2_. The culture medium was removed and replaced with fresh medium in the presence of (i) 25 μM azithromycin (MCE, Monmouth Junction, NJ, USA), (ii) 50 μM azithromycin, and (iii) 300 μM Zn (Sigma-Aldrich, ST. Louis, MO, USA), (iv) 300 μM Zn and 25 μM azithromycin, (v) 300 μM Zn and 50 μM azithromycin, and incubated for 24 or 48 hours.

### RNA isolation

Total RNA was isolated using the PureLink™ RNA mini kit (Thermo Fisher Scientific, Waltham, MA, USA) according to the manufacturer’s instructions. The RNA concentration and quality were assessed using the Quibit 3.0 Fluorometer (Thermo Fisher Scientific). Total RNA samples were stored at −80 °C.

### Reverse transcription (RT)

RT reactions were carried out using the superscript III first strand synthesis system (Thermo Fisher Scientific, Waltham, MA, USA). cDNA was synthesized starting from 2 μg of purified total RNA. The reactions in a final volume of 20 μl contained 1x Buffer, DTT, dNTPs, superscript III RT, and 500 ng oligo(dT). Samples were incubated at 65 °C for 5 minutes and 50 °C for 60 minutes, and then the RT enzyme was inactivated by heating to 70 °C for 15 minutes. cDNA samples were stored at −20 °C.

### Quantitative PCR (qPCR)

qPCR was carried out with KAPA SYBR^®^FAST qPCR Master Mix (Wilmington, MA, USA) in a final volume of 20 μl, with 0.3 μM forward and reverse primer and 1 μl of cDNA. Fluorescent detection was performed using the ABI 7500 fast System (Thermo Fisher Scientific) with the following thermal cycling conditions: initial polymerase activation at 95 °C for 3 minutes, followed by 40 cycles of denaturation at 95 °C for 3 seconds and annealing/extension at 60 °C for 30 seconds. After amplification, dissociation (melting) curve analysis was performed to analyze the product melting temperature. Each sample was amplified in triplicate wells. Negative (no template) controls were included in each assay. The results were analyzed using ABI 7500 Fast System Software. The threshold cycle (Ct) at which the amount of amplified target reached a fixed threshold was determined. Relative expression was calculated using the 2^−ΔΔCq^ method. The results were analyzed and are shown as the fold change relative to each control group.

**Table 1.**
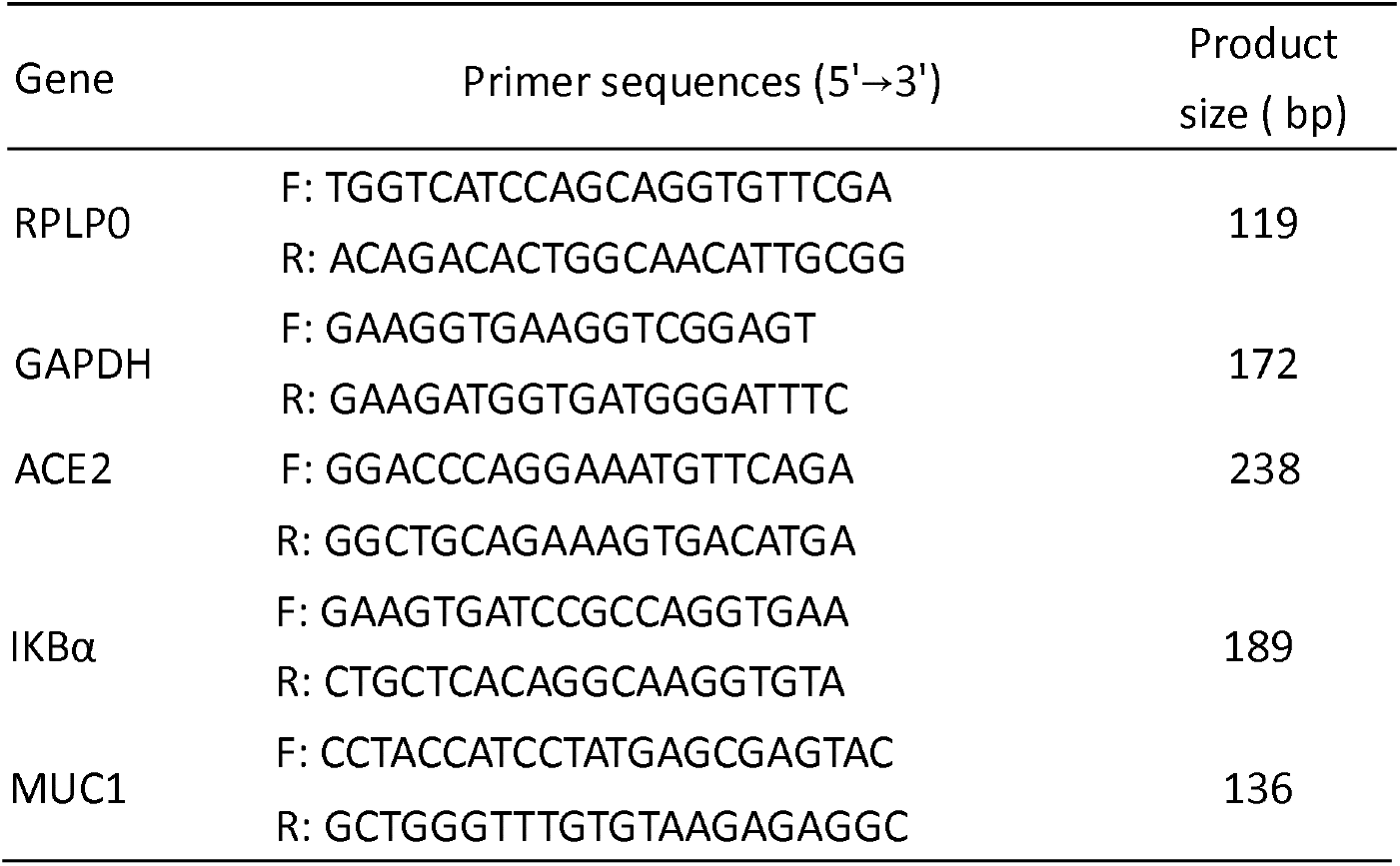
Primers used for aPCR.

### Western blot

Cells were scraped with lysis buffer (1% Triton X-100, 20◻mM Tris pH 7.4, 150◻mM NaCl, and protease inhibitors) on an ice tray, and cell lysates were subjected to western blot analysis. Protein samples were first separated by SDS–PAGE and then transferred to a PVDF membrane. Primary antibodies were applied to detect specific protein expression, followed by incubation with appropriate HRP-conjugated secondary antibodies. Protein signals were developed using an enhanced chemiluminescence reagent (Biomate, Taiwan) and detected by a BIO-RAD ChemiDoc™ MP imaging system (BIO-RAD, Hercules, CA, USA). Western blots were carried out with ACE-2 antibody (Bioss, Woburn, MA, USA), IKB-α antibody (Santa Cruz, Dallas, TX, USA) and PARP antibody (Cell Signaling, Danvers, MA, USA); α-tubulin antibody (Novus, Littleton, CO, USA) was used as a loading control.

### Statistical analysis

For statistical analysis, the mean and standard errors were calculated by using GraphPad Prism software version 9 (GraphPad Software Inc., San Diego, CA, USA). Student’s t-tests were used to determine significant differences between two experimental conditions. Data were presented as mean + SEM.

## Author contributions

YKC, TTH, MCL and BRL conceived and designed the research; CWC and YJH performed the experiments; CWC and YKC wrote the manuscript and generated the illustrations. TTH, YKC, BRL, YPL and CYC reviewed and edited the manuscript; All authors read and approved the final manuscript.

## Acknowledgments

We are grateful to Ann-Lii Cheng (Dean of National Taiwan University Cancer Center) and Dr. Ming-Hua Shiao, Dr. Yao-Joe Joseph Yang (Chiefs of Taiwan Instrument Research Institute, National Applied Research Laboratories) for organization and coordination between the groups. We thank staff of the Second Core Lab, Department of Medical Research, National Taiwan University Hospital for technical support during the study. We thank Chao-Chi Ho (National Taiwan University Hospital) for technical support of lung cancer cell lines.

